# Excitation-contraction coupling plasticity is revealed upon RyR2 depletion in adult mouse ventricular myocytes

**DOI:** 10.64898/2026.10.07.755981

**Authors:** Erick B. Ríos Pérez, Achal M. Gowda, Isabel C. Garay, Sergio Estrada, David W. Kingston, Francisco J. Alvarado

## Abstract

Ryanodine receptor 2 (RyR2) is central to cardiac excitation-contraction coupling (ECC), translating electrical signals into sarcoplasmic reticulum (SR) Ca^2+^ release and mechanical contraction. Although constitutive deletion of RyR2 is lethal, adult hearts can withstand a significant reduction in RyR2 expression without overt ECC dysfunction. We hypothesized that ECC possesses intrinsic homeostatic plasticity that rapidly adapts to acute RyR2 loss to maintain cardiac performance. Using an adult tamoxifen-inducible RyR2 conditional knockout (RyR2cKO) mouse model, we evaluated cardiac structure, electrophysiology, and intracellular Ca^2+^ dynamics. RyR2cKO mice survived for weeks with <10% residual RyR2 protein expression. In vivo echocardiography and electrocardiography revealed normal cardiac structure and function. Confocal Ca^2+^ imaging in intact hearts and isolated ventricular myocytes showed preserved Ca^2+^ transient (CaT) amplitude but slower release kinetics and elevated SR load. Simultaneous patch-clamp and confocal Ca^2+^ imaging established that L-type Ca^2+^ current (*I*_Ca_) and ECC gain were maintained in RyR2cKO myocytes. Expression of Ca_V_1.2, NCX, and phospholamban remained unchanged, whereas SERCA2a, IP_3_ receptor, and VMP1 levels decreased. Pharmacological inhibition of IP_3_ receptors did not suppress CaT, ruling out alternative SR channel compensation. These findings reveal remarkable intrinsic ECC plasticity, where high RyR2 channel reserve, elevated SR Ca^2+^ load, and CaT autoregulation collectively maintain cardiac Ca^2+^ cycling and contractile function despite profound RyR2 depletion.

**KEY POINTS:**

- Deletion of RyR2, a critical channel for cardiac excitation-contraction coupling (ECC), is embryonically lethal; however, adult cardiomyocytes can tolerate a significant reduction in RyR2 expression without overt dysfunction.
- We show that mouse hearts and cardiomyocytes with a 90% reduction in RyR2 expression show preserved Ca^2+^ handling.
- Compensation for reduced RyR2 levels does not involve overt remodeling of the ECC machinery or cardiac structure.
- Cardiomyocytes have a RyR2 reserve and employ autoregulation to maintain ECC fidelity, uncovering homeostatic plasticity as a regulatory mechanism of ECC.
- Reduced RyR2 expression is often observed in heart disease; however, it may not contribute to the pathology.

## INTRODUCTION

Excitation-contraction coupling (ECC) is the process that converts electrical activity into mechanical contraction in the heart. Excitation is driven by the action potential (AP), generated by highly choreographed opening of voltage-gated Na^+^, Ca^2+^ and K^+^ channels. Ca^2+^ influx through voltage-gated L-type Ca^2+^ channels (LTCCs, *I*_CaL_) during the AP activates ryanodine receptor 2 (RyR2) channels through Ca^2+^-induced Ca^2+^-release (CICR). Ca^2+^ released by RyR2 from the sarcoplasmic reticulum (SR), an intracellular store, floods the cytosol and triggers contraction of the myofilaments. Relaxation occurs when Ca^2+^ is pumped back into the SR by the SR Ca^2+^ ATPase 2 (SERCA2) or extruded from the cytosol by the Na^+^/Ca^2+^ exchanger (NCX) (Bers, 2002).

Constitutive knockout (KO) of most ECC proteins is embryonically lethal (Takeshima *et al*., 1998; Periasamy *et al*., 1999; Seisenberger *et al*., 2000; Reuter *et al*., 2002); however, many studies have shown that their expression can be significantly reduced without severely compromising ECC or cardiac function. Heterozygous KO of RyR2 in mice and rabbits leads to a ∼50% decrease without inducing dysregulation of Ca^2+^ handling (Zou *et al*., 2011; Zheng *et al*., 2022). A ∼95% reduction in SERCA2 expression in a conditional KO mouse model was compensated for by enhanced *I*_CaL_ and NCX activity to preserve quasi-normal heart function (Andersson *et al*., 2009). Moreover, a cardiac-specific model with an ∼80% decrease in NCX expression showed remarkable adaptation to preserve function by limiting *I*_CaL_ and increasing reliance on intracellular Ca^2+^ cycling via RyR2 and SERCA2 (Henderson *et al*., 2004; Lotteau *et al*., 2021). Altogether, these studies support the idea that compensatory mechanisms are in place to regulate both ECC and excitability and maintain function when the expression of key proteins is altered. In the nervous system, homeostatic plasticity describes the ability of neurons to stabilize activity through compensatory adjustments in protein function and expression (Turrigiano, 2012). This process ensures that firing rates and synaptic transmission are preserved under conditions of altered excitability. Although extensively characterized in the nervous system, analogous mechanisms have not been formally demonstrated in cardiomyocytes. Nevertheless, emerging evidence suggests that plasticity in the heart can be achieved through multi-layered coordination of ion channel biogenesis and spatial coordination (Ponce-Balbuena *et al*., 2018; Jameson *et al*., 2023).

We hypothesized that homeostatic plasticity contributes to maintaining Ca^2+^ handling and cardiac function when the expression of ECC proteins is reduced. To explore this possibility, we employed a tamoxifen (TAM)-inducible conditional RyR2 knockout mouse model (RyR2cKO), as previously reported (Bround *et al*., 2012; Bround *et al*., 2016). Post-TAM injection, RyR2 expression progressively decreases in RyR2cKO hearts eventually leading to death (Bround *et al*., 2012). Using this model, we sought to evaluate the properties of Ca^2+^ handling and ECC integrity at specific time-points with a defined level of remnant RyR2. We demonstrate that ECC is a highly robust process, capable of quickly adapting to a dramatic loss of RyR2 to preserve heart function. We show that mice can survive with less than 10% of RyR2 while Ca^2+^ handling is largely preserved in whole hearts and isolated cardiomyocytes, and cardiac function is normal in intact animals. Altogether, our results support the hypothesis that homeostatic cardiac plasticity occurs in ventricular myocytes to maintain ECC fidelity.

## MATERIALS AND METHODS

### Ethical approval

Animal studies were approved by the University of Wisconsin-Madison School of Medicine and Public Health Institutional Animal Care and Use Committee (Protocol M005944), accredited by the Association for Assessment and Accreditation of Laboratory Animal Care International. Husbandry was performed by specialized personnel of the School of Medicine and Public Health’s Biomedical Research Model Services using standard procedures. Animals were kept in specialized rooms with humidity and temperature control in a 12 h light/dark cycle. Animal care was provided by veterinarians of the Research Animal Program and Compliance.

### Reagents

All reagents were obtained from MilliporeSigma unless otherwise stated. A list of catalog numbers and commercial vendors is available upon request.

### RyR2 conditional knockout mouse model

Floxed RyR2 mice (Ryr2^flox/flox^) (JAX stock #026628) (Bround *et al*., 2012) and αMHC-MerCreMer mice (JAX stock #005657) (Sohal *et al*., 2001) were obtained from Jackson Laboratories and crossed to obtain a cardiac-specific, TAM-inducible RyR2 knockout mouse model, as previously described (Bround *et al*., 2012). Experimental RyR2cKO mice were Ryr2^flox/flox^ and hemizygous for the αMHC-MerCreMer transgene, while controls were Ryr2^flox/flox^ and MerCreMer negative (**Figure 1A**). Experimental and control mice of 3 to 6 months of age received intraperitoneal injections of TAM (0.1 mg/g dissolved in peanut oil with 10% ethanol (Cerrone *et al*., 2014)) for three consecutive days (Bround *et al*., 2012). Mice were used for experiments one and three weeks post-injections (counted from the day of the first injection). All experiments included male and female mice, and animals were age matched.

**Figure 1.**
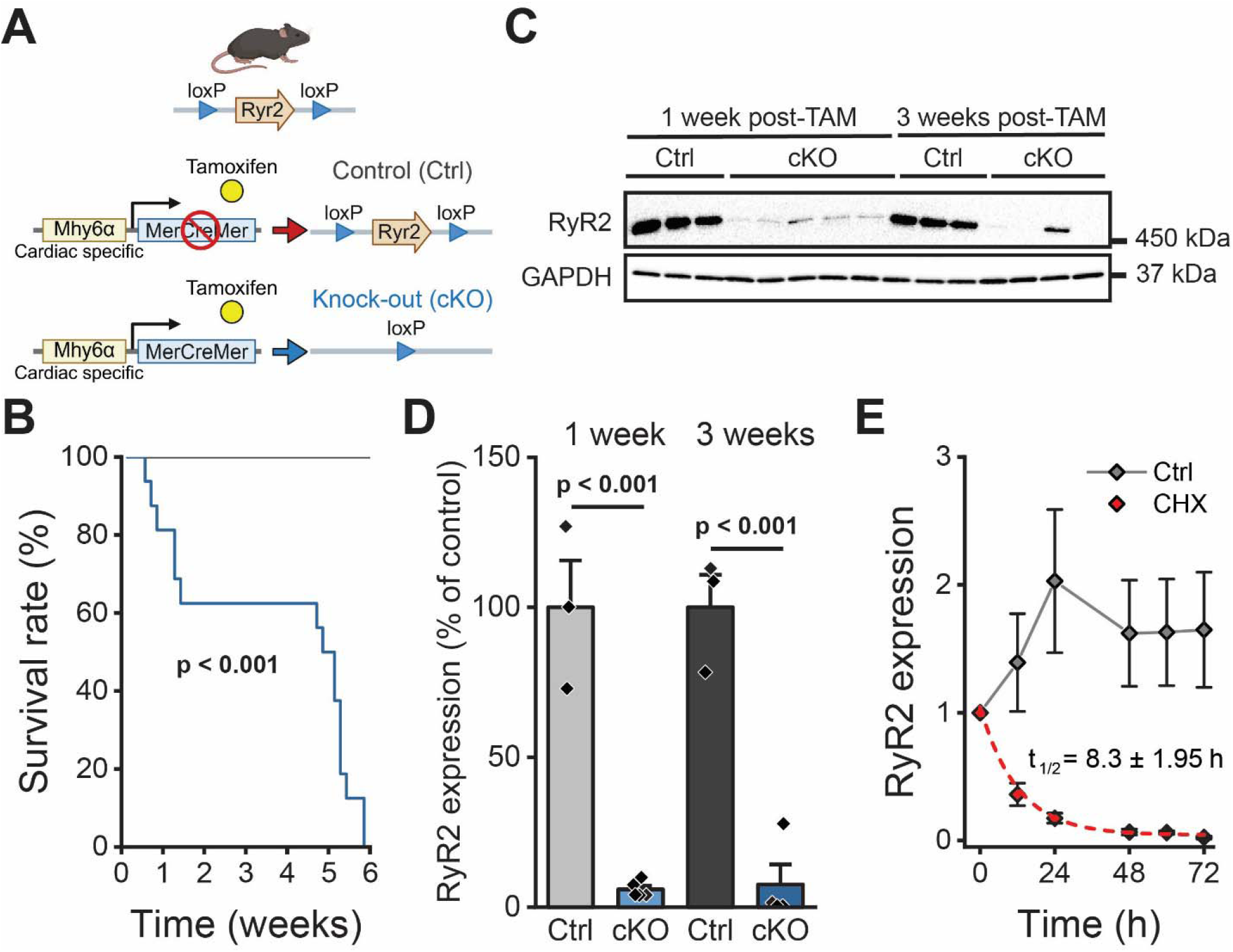
RyR2 expression is significantly reduced within 1-week of tamoxifen treatment in RyR2cKO hearts. **A**) Diagram of the RyR2cKO mouse model and experimental groups used for the study. **B**) Survival plot of control (gray) and RyR2cKO (blue) mice after tamoxifen (TAM) treatment. N = 16 for both groups. Log-rank (Mantel-Cox) test. **C**) Representative blots of RyR2 expression from heart homogenates of control (Ctrl) and RyR2cKO (cKO) mice collected 1- and 3-weeks post-TAM. **D**) Quantification of RyR2 expression in control and RyR2cKO heart homogenates. N = 3 for Ctrl and 4 for cKO. **E**) Cycloheximide (CHX) chase assay in a HEK293 cell-line stably expressing RyR2. After RyR2 expression was induced for 48 h, protein translation was blocked using 75 µM of CHX or vehicle (Ctrl) and cells were collected at different times to measure RyR2 expression using Western blots. RyR2 expression was normalized to the signal at time zero. N = 4 independent experiments per group. Data were fitted to an exponential decay function *y* = *y*_0_ + Ae^-x/t^, where *y* represents RyR2 expression. Linear mixed-effects model (LMM) with Bonferroni post-hoc. Panel A was created in BioRender https://biorender.com/h6tny7r.

### Tissue homogenization

Heart homogenates were prepared as established previously (Zheng *et al*., 2025). Briefly, explanted hearts were flash-frozen in liquid nitrogen and stored at -80°C until used. Hearts were pulverized in liquid nitrogen, suspended in homogenization buffer (0.9% NaCl, 10 mM Tris-HCl (pH 6.8), 20 mM NaF, 2 µM leupeptin, 100 µM phenylmethylsulfonyl fluoride, 500 µM benzamidine, and 100 nM aprotinin), homogenized using a Teflon pestle and centrifuged at 1,000 x *g* for 8 min at 4°C. Supernatants were aliquoted and stored at -80°C until further use. Protein concentrations were measured using the Bradford method (Bio-Rad).

### Western blot assays

50 µg of tissue homogenate or cell lysate were suspended in Laemmli buffer and separated by SDS-PAGE in 4-20% TGX precast gels (Bio-Rad). Proteins were transferred to PVDF membranes using iBlot2 transfer system (ThermoFisher) or overnight wet transfer. Membranes were then probed using the following antibodies: RyR (clone 34C, 1:2,000; ThermoFisher, MA3-925), SERCA2 (clone 2A7-1, 1:1,000; Abcam, ab2861), NCX1 (clone EPR12739, 1:1,000; Abcam, ab177952), Ca_V_1.2 (1:200; Alomone, ACC-003), PLB (clone A1, 1:5,000; Badrilla, A010-14), VMP1 (clone D1Y3E, 1:1,000; Cell Signaling, 12929), IP_3_R (clone A-5, 1:250, Santa Cruz Biotechnology, SC-398434), vinculin (clone EPR19579, 1:1,000; Abcam, ab76083) and GAPDH (clone 6C5, 1:10,000; Millipore, MAB374). Secondary antibodies, used as appropriate, included goat anti-mouse-HRP (1:1,000 or 1:5,000; Thermo Fisher Scientific, 31437) or goat anti-rabbit-HRP (1:1,000 or 1:5,000; Thermo Fisher Scientific, 31463). Membranes were developed with Clarity ECL reagent (Bio-Rad) or SuperSignal ECL reagent (ThermoFisher) and imaged with the ChemiDoc MP system (Bio-Rad). Band intensities were quantified using ImageLab 6.1.0 software (Bio-Rad).

### Cycloheximide chase assay

Stable tetracycline-inducible HEK293 cell lines expressing WT-RyR2 (Alvarado *et al*., 2019) were grown in high-glucose Dulbecco’s Modified Eagle Medium (DMEM, Corning) supplemented with 10% fetal bovine serum (FBS, ThermoFisher), 0.1mM non-essential amino acids (ThermoFisher), 1mM sodium pyruvate (ThermoFisher), and 2mM GlutaMAX (ThermoFisher) on 100-mm tissue culture dishes. RyR2 expression was induced by adding tetracycline (1 µg/mL). 48 hours post-tetracycline induction, cells were washed with PBS, and culture medium was replaced with fresh medium containing 75 μM cycloheximide (CHX) to inhibit protein translation. Control plates were maintained in parallel without addition of CHX to the culture medium. Cells were harvested at 0-, 12-, 24-, 48-, 60-, and 72-hours post-treatment. Cells were solubilized in lysis buffer containing 25 mM Tris/50 mM HEPES (pH 7.4), 137 mM NaCl, 1% CHAPS, 0.5% soybean phosphatidylcholine, 2.5 mM DTT and protease inhibitors. After incubation on ice for 1 hour, the lysates were centrifuged at 10,000 x *g* at 4 °C for 10 min and the supernatants were collected for further processing. Protein concentrations were measured using the Bradford method (Bio-Rad) and Western blots were performed as described above.

### Echocardiography

Transthoracic echocardiography was conducted by the Small Animal Imaging and Radiotherapy Facility (SAIRF) at the University of Wisconsin-Madison using a Vevo 3100 system running Vevo Lab 5.8.2 (Visual Sonics) with a 22 – 55 MHz transducer (Visual Sonics, MS550D). Mice were anesthetized with 1 – 2% isoflurane and maintained on a heated platform. The temperature was monitored and recordings were performed at 37.0 ± 0.5 °C. 2D M-mode images from the left ventricle in the long axis were acquired. Left ventricle diameter, septum and posterior wall thickness were recorded to calculate heart function parameters (fractional shortening, left ventricle volume, ejection fraction, heart rate, cardiac output, etc.). All parameters were obtained using VevoLab software and measured after at least the first three consecutive cycles. Imaging was conducted in a single-blind manner to avoid biases from the echocardiography technician.

### Electrocardiography

Surface electrocardiograms were recorded as previously described (Zheng *et al*., 2025). Mice were anesthetized using 2% isoflurane and kept on top of a heated pad to maintain the temperature at 37.0 ± 0.5 °C. Needle electrodes were placed under the skin to record in Lead-I and Lead-II configurations, using a PowerLab 8/35 system and LabChart 8.1.21 (ADInstruments). After the mice reached a stable temperature, baseline recordings were obtained for at least 10 minutes. ECGs were analyzed using LabChart 8.1.21 software.

### Tissue immunostaining

Mouse heart tissue samples were fixed in 10% formalin (Buffered Formalde-Fresh SF93-4, Fisher Scientific) for at least 24 h and then maintained in 70% ethanol before being processed by the Translational Research Initiatives in Pathology (TRIP) laboratory of the University of Wisconsin-Madison. Briefly, samples were incubated overnight with a series of ethanol, xylenes, and paraffin. Mouse heart tissue samples were embedded in formalin-fixed paraffin-embedded (FFPE) blocks. 5 µm tissue sections were deparaffinized, and antigens were retrieved in boiling 1X Antigen Retrieval buffer pH 9.0 for 8 min. Tissue sections were blocked with 1X Diluent/Block (Akoya). Tissues were then washed and stained at room temperature with anti-RyR2 (Abcam, ab302716) and anti-N-Cadherin (Abcam, ab76011). Tissues were washed and stained with secondary antibodies using goat anti-rabbit (Abcam, ab6721), followed by washing and staining with Opal dyes 650 and 520, respectively. This process was sequentially repeated for each antibody. Tissues were washed and stained with DAPI Working Solution (Akoya) before mounting in ProLong™ Gold Anti-fade mounting medium (ThermoFisher). Immunostaining was performed in a single-blind manner to avoid biases from the technician. Images were collected using a ZEISS LSM 980 Airyscan 2 system using a 63x/1.40 oil immersion objective. Image acquisition parameters were optimized using ZEN 3.14 software (ZEISS).

### Ca^2+^ transient recordings in intact hearts

Ca^2+^ transients (CaTs) in intact hearts were recorded as previously published (Wang *et al*., 2025). Excised hearts were mounted on a Langendorff system and perfused with a modified Krebs-Henseleit (KB) buffer (in mM; NaCl 120, NaHCO_3_ 24, Glucose 11.1, KCl 5.4, MgCl_2_ 1, NaH_2_PO_4_ 0.42, taurine 10, creatine 5, pH 7.4) to remove blood from the heart. Hearts were then loaded with 10 µM Rhod-2 AM (ThermoFisher) in KB buffer for 30 min at room temperature. Hearts were placed in a recording chamber mounted onto the stage of a confocal Zeiss LSM800 microscope for in-situ imaging. Perfusion was maintained using KB buffer supplemented with 1.8 mM CaCl_2_ and 10 µM blebbistatin (MedChemExpress) to prevent motion artifacts. Confocal line scans were recorded for 30 s on hearts paced at 6 Hz using a 40X/1.2 water immersion objective at 1.82 ms per line. Rhod-2 was excited at 553 nm and emission was recorded at 577 nm. Data analysis was performed using ZEN 2.6 (ZEISS) and custom MATLAB 2026a scripts were used to obtain CaT parameters (MathWorks). To obtain CaT parameters, F_0_ was established as the fluorescence signal at the end of a CaT instead of using a region with no signal.

### Isolation of ventricular myocytes

Isolated ventricular myocytes were obtained using a Langendorff perfusion system as reported previously (Gowda *et al*., 2026). Mice were injected with heparin (0.5 U/g, intraperitoneal) prior to dissection of the heart. Cervical dislocation was used to sacrifice the mice. The heart was then quickly excised, cannulated, mounted on a Langendorff system and perfused with perfusion buffer containing (in mM): 113 NaCl, 4.7 KCl, 1.2 MgSO_4_ · 7 H2O, 10 HEPES, 0.6 Na_2_HPO4, 12 NaHCO_3_, 0.6 KH_2_PO_4_, 10 KHCO_3_, 30 taurine, 500 2,3-butanedione monoxime and 5.5 glucose (pH 7.40) at 37 °C at 3 ml/min. After most of the blood was cleared from the heart, perfusion buffer was switched to a digestion buffer (perfusion buffer supplemented with 775 U/ml collagenase type II (Worthington), and 12.5 µM CaCl_2_). The heart was digested for 7 – 10 min, after which the ventricles were cut, and cells were mechanically dissociated from the tissue in stopping buffer (perfusion buffer plus 12.5 µM CaCl_2_ and 10% fetal bovine serum [FBS]). Ca^2+^ was then slowly reintroduced to the cell suspension in six steps until reaching a 1.8 mM Ca^2+^ concentration. Cells were then transferred to Tyrode’s solution containing (in mM): 135 NaCl, 4 KCl, 1.8 CaCl_2_, 1 MgCl_2_, 10 HEPES, 1.2 NaH_2_PO_4_ and 10 glucose (pH 7.40) until used.

### Ca^2+^ transients

Ca^2+^ transients were recorded as previously described (Gowda *et al*., 2026). To obtain Ca^2+^ transients and SR load, isolated cardiomyocytes were loaded with 10 µM fluo-4 AM in 0.4% pluronic F-127 (ThermoFisher) for 5 minutes at 37°C. Cells were then plated on laminin-coated glass-bottom dishes (MatTek) with normal Tyrode’s solution and imaged using a LSM 980 confocal microscope with a 40x/1.2 water immersion objective at 1.23 ms per line using ZEN 3.13 software (Zeiss). A 488 nM excitation laser was used and emission was recorded at 516 nm. Longitudinal line scans were recorded during 1, 2, and 3 Hz field stimulation for 30, 15, and 15 s, respectively (2-ms pulses, 60–80 V). A 5 s pulse of 10 mM caffeine was applied at the end of the 3 Hz stimulation to obtain SR load. For measuring CaT parameters, the last 10 transients from each stimulation frequency were averaged. Recordings were obtained in normal Tyrode’s solution at room temperature. CaT properties were obtained using a custom MATLAB script and run using MATLAB 2026a (MathWorks) (Zheng *et al*., 2025).

### Simultaneous patch-clamp and Ca^2+^ imaging

Patch-clamp experiments were performed using the whole-cell configuration with ruptured patch, as reported elsewhere (Janicek *et al*., 2024). Freshly isolated cells were resuspended in Tyrode’s solution supplemented with 5 mM 4-aminopyridine (4-AP) and 30 µM tetrodotoxin (TTX, Cayman Chemical) to block *I*_to_ and *I*_Na_, respectively. Recording pipettes were filled with an internal solution containing (in mM): 110 CsCl, 6 MgCl_2_, 5 Na_2_ATP, 0.3 Na_2_GTP, 10 HEPES and 15 TEA·Cl, pH adjusted to 7.2 with CsOH and supplemented with 200 µM fluo-4 pentapotassium salt to record Ca^2+^ signals. Once filled, the pipette resistance was 2 – 5 MΩ. Upon patch rupture, cells were dialyzed to allow fluo-4 equilibration until a stable fluorescent signal was achieved. *I*_CaL_ was measured by holding the cells at -40 mV and applying 300 ms voltage steps from -40 to 50 mV in 10 mV increments with 2 s intervals between each sweep. Simultaneously, CaTs were recorded using confocal microscopy as described above.

### Statistical analysis and reproducibility

Control and RyR2cKO mice were independently treated with TAM for each set of experiments and data were collected at specific timepoints for both experimental groups. Data are presented as the mean ± s.e.m. with individual data points overlaid, unless otherwise specified. Statistical significance was determined at p < 0.05. Sample sizes are indicated in each figure legend. Comparisons between control and RyR2cKO mice or cells were done using a Linear mixed-effects model (LMM) with Bonferroni post hoc analysis. LMM or two-tailed t-tests were used to compare protein expression levels between control and cKO samples. Statistical tests for each experiment are specified in all figure legends. All statistical tests were performed using IBM SPSS software version 32 or SigmaPlot 15 (Grafiti LLC). Data were managed in Excel or Origin, and graphs were generated in Origin 2025.

## RESULTS

### RyR2-conditional knockout mice survive with remarkably low RyR2 expression levels

We used a cardiac-specific, tamoxifen (TAM)-inducible RyR2cKO model previously reported (Bround *et al*., 2012). We first determined the survival rate of mice after TAM injections to identify the best time points to conduct this study. Remarkably, RyR2cKO mice survived for up to 6 weeks (**Figure 1B**). Within one week post-TAM injections there was already a ∼90% reduction in RyR2 expression in RyR2cKO hearts (**Figure 1C,D**). At three weeks post-TAM, three out of four RyR2cKO hearts showed almost no detectable RyR2 expression as determined by Western blot (**Figure 1C,D**) and the mean expression level was below 10% of that of control mice. Previous studies have demonstrated that the MerCreMer transgene can induce transient cardiomyopathy during the first days after TAM injections, potentially leading to sudden death (Koitabashi *et al*., 2009). Hence, the high mortality observed in RyR2cKO mice in the first week may be associated with genetic recombination and MerCreMer activation. Similar transient cardiomyopathy was reported previously in RyR2cKO mice (Bround *et al*., 2012). Surprisingly, RyR2cKO mice that survive the first week post-TAM enter a stable period in which they are indistinguishable from the control group. Altogether, we found that decreasing RyR2 expression by more than 90% does not cause immediate death, suggesting that ECC is a robust process.

Given the fast decrease in RyR2 expression in RyR2cKO hearts, we performed a cycloheximide (CHX) chase assay in a RyR2 inducible HEK cell line to estimate the RyR2 turnover rate. We found that upon CHX application, the RyR2 protein degrades within 24 h with a half-life of ∼8 hours (**Figure 1E**). This suggests that fast degradation kinetics can lead to an abrupt decrease in RyR2 expression in RyR2cKO myocytes.

### RyR2 knockout does not affect overall heart structure or function

We investigated overall heart function and structure in a cohort of control and RyR2cKO mice using electro and echocardiography one and three weeks post-TAM. Mice that died after the first assessment (one week) were excluded from this dataset. Electrocardiography measurements showed an increase in the QTc duration in RyR2cKO mice at one week post-TAM, but such a difference was not present at three-weeks (**Table 1**). In RyR2cKO mice at three weeks post-TAM we only observed an increase in the QRS interval duration, while all the other parameters were comparable to those of control mice (**Table 1**). We further analyzed heart structure using echocardiography (**Figure 2A,B and Table 2**). No differences in heart rate, ejection fraction, fractional shortening, or left ventricle diameter were observed between control and RyR2cKO mice (**Figure 2C-F**). We found a transient decrease in the global longitudinal strain (GLS) in RyR2cKO mice at one-week post-TAM that was no longer present at three weeks (**Table 2**). Altogether, these results show that a significant decrease of ∼90% in RyR2 expression does not overtly affect cardiac structure and function, suggesting the presence of a compensatory network that preserves ECC function. The transitory changes we observed at one-week post-TAM suggest an effect that may be associated with the induction of the MerCreMer transgene model (Koitabashi *et al*., 2009) in addition to the dramatic decrease in RyR2 expression. For this reason, we conducted further studies only in mice that survived 3-weeks post-TAM.

**Figure 2.**
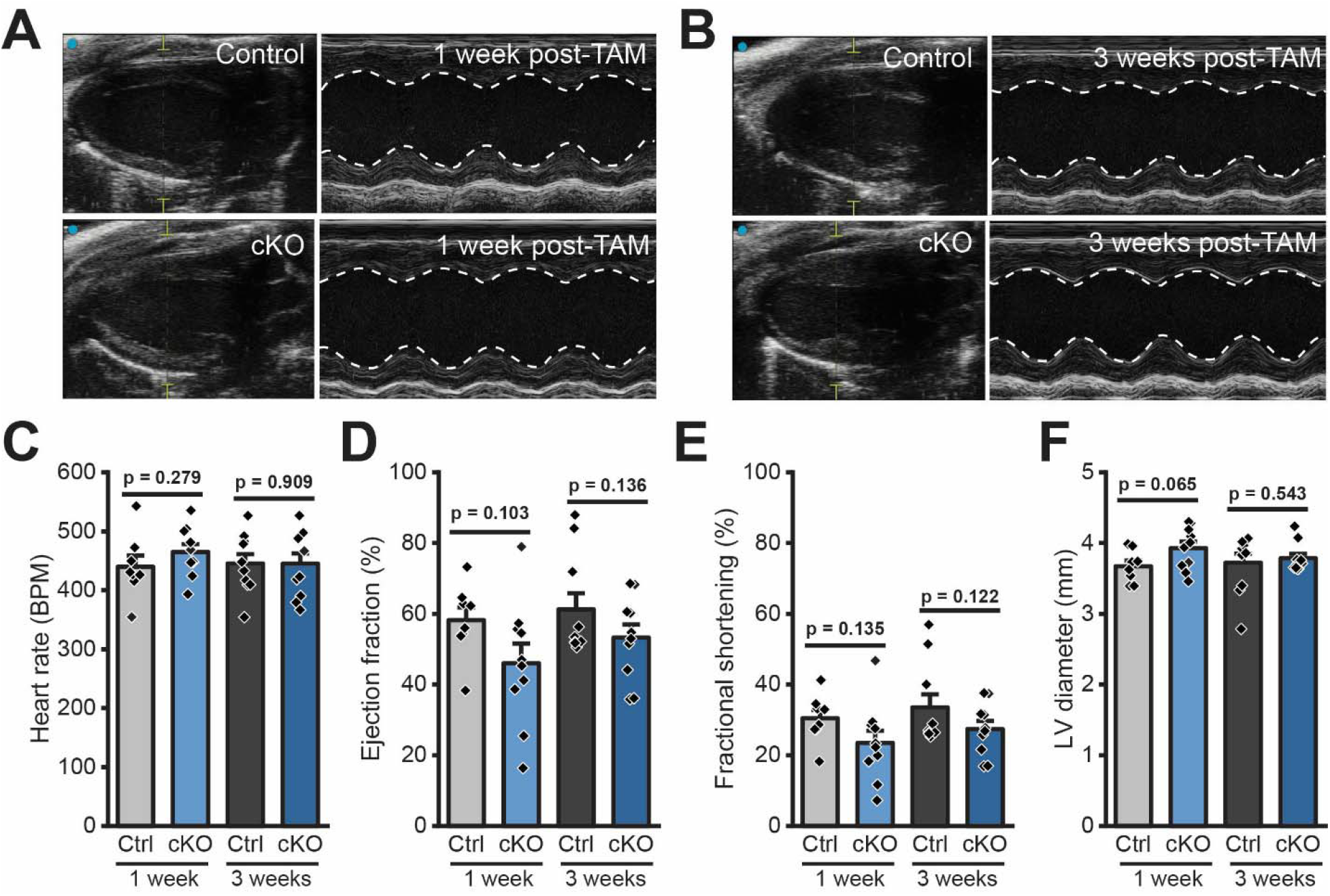
Cardiac function and structure are preserved in RyR2cKO hearts. **A,B**) Representative M-mode echocardiography images recorded from control (top) and RyR2cKO (bottom) 1 and 3 weeks post-TAM injection. **C-F**) Mean heart rate (C), ejection fraction (D), fractional shortening (E) and systolic LV diameter (F) in control (Ctrl) and RyR2cKO (cKO) mice at 1- and 3-weeks post-TAM. N = 8 Ctrl and 10 cKO mice. Linear mixed-effects model (LMM) with Bonferroni post-hoc.

**Table 1.** Electrocardiography parameters in control and cKO mice under anesthesia.

|  | 1-week post-TAM |  | 3-weeks post-TAM |  |
| --- | --- | --- | --- | --- |
|  | Ctrl (N = 12) | cKO (N = 8) | Ctrl (N = 12) | cKO (N = 8) |
| Heart rate (BPM) | 522.7 ± 13.1 | 518.8 ± 12.7 | 524.5 ± 13.2 | 500.9 ± 16.0 |
| RR interval (ms) | 115.7 ± 2.9 | 116.2 ± 2.8 | 115.3 ± 3.1 | 120.8 ± 4.0 |
| PR interval (ms) | 39.2 ± 1.0 | 38.2 ± 1.1 | 39.5 ± 1.1 | 38.8 ± 0.7 |
| P wave duration (ms) | 21.7 ± 1.6 | 21.8 ± 2.4 | 24.2 ± 1.4 | 23.6 ± 1.8 |
| QRS interval (ms) | 7.9 ± 0.4 | 7.8 ± 0.4 | 7.8 ± 0.7 | 10.2 ± 0.9<br>* (p = 0.044) # (p = 0.005) |
| QTc (ms) | 58.5 ± 2.7 | 72.2 ± 4.9<br>* (p = 0.025) | 59.7 ± 3.7 | 62.6 ± 4.6 |

\* Comparison between groups at the same time points.

**Table 2.**
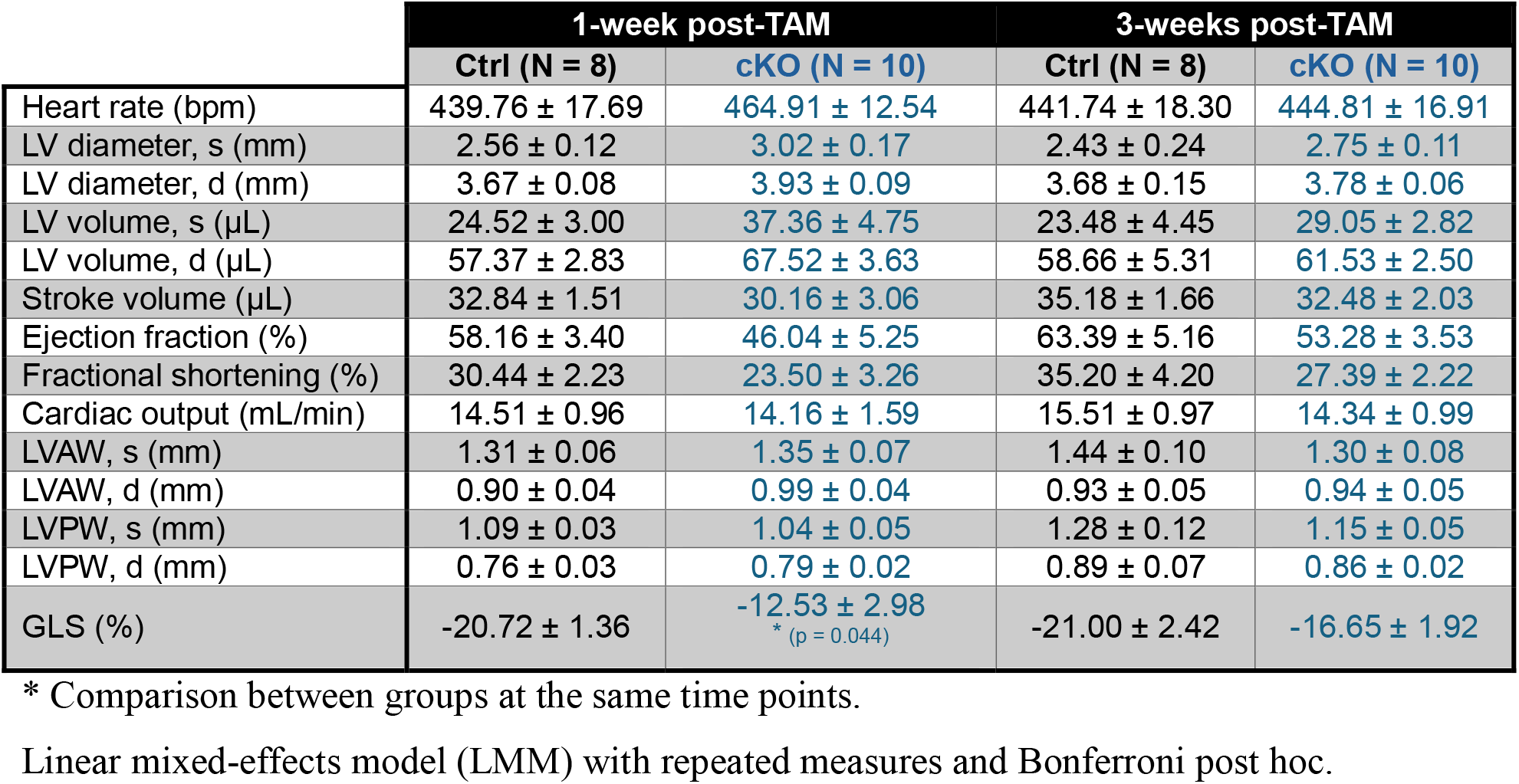
Echocardiography parameters in control and cKO mice under anesthesia.

|  | 1-week post-TAM |  | 3-weeks post-TAM |  |
| --- | --- | --- | --- | --- |
|  | Ctrl (N = 8) | cKO (N = 10) | Ctrl (N = 8) | cKO (N = 10) |
| Heart rate (bpm) | 439.76 ± 17.69 | 464.91 ± 12.54 | 441.74 ± 18.30 | 444.81 ± 16.91 |
| LV diameter, s (mm) | 2.56 ± 0.12 | 3.02 ± 0.17 | 2.43 ± 0.24 | 2.75 ± 0.11 |
| LV diameter, d (mm) | 3.67 ± 0.08 | 3.93 ± 0.09 | 3.68 ± 0.15 | 3.78 ± 0.06 |
| LV volume, s (μL) | 24.52 ± 3.00 | 37.36 ± 4.75 | 23.48 ± 4.45 | 29.05 ± 2.82 |
| LV volume, d (μL) | 57.37 ± 2.83 | 67.52 ± 3.63 | 58.66 ± 5.31 | 61.53 ± 2.50 |
| Stroke volume (μL) | 32.84 ± 1.51 | 30.16 ± 3.06 | 35.18 ± 1.66 | 32.48 ± 2.03 |
| Ejection fraction (%) | 58.16 ± 3.40 | 46.04 ± 5.25 | 63.39 ± 5.16 | 53.28 ± 3.53 |
| Fractional shortening (%) | 30.44 ± 2.23 | 23.50 ± 3.26 | 35.20 ± 4.20 | 27.39 ± 2.22 |
| Cardiac output (mL/min) | 14.51 ± 0.96 | 14.16 ± 1.59 | 15.51 ± 0.97 | 14.34 ± 0.99 |
| LVAW, s (mm) | 1.31 ± 0.06 | 1.35 ± 0.07 | 1.44 ± 0.10 | 1.30 ± 0.08 |
| LVAW, d (mm) | 0.90 ± 0.04 | 0.99 ± 0.04 | 0.93 ± 0.05 | 0.94 ± 0.05 |
| LVPW, s (mm) | 1.09 ± 0.03 | 1.04 ± 0.05 | 1.28 ± 0.12 | 1.15 ± 0.05 |
| LVPW, d (mm) | 0.76 ± 0.03 | 0.79 ± 0.02 | 0.89 ± 0.07 | 0.86 ± 0.02 |
| GLS (%) | -20.72 ± 1.36 | -12.53 ± 2.98<br>* (p = 0.044) | -21.00 ± 2.42 | -16.65 ± 1.92 |

\* Comparison between groups at the same time points.
Linear mixed-effects model (LMM) with repeated measures and Bonferroni post hoc.

### Whole-heart Ca^2+^ transients are indistinguishable between control and RyR2 knockout hearts

To investigate how intracellular Ca^2+^ dynamics were affected by RyR2 expression levels, we performed in-situ Ca^2+^ imaging in intact RyR2cKO and control hearts at three weeks post-TAM injection. Hearts were loaded with Rhod-2 and imaged under a confocal microscope while electrical pacing was applied at 6 Hz (Wang *et al*., 2025). Representative traces from control and RyR2cKO hearts show that Ca^2+^ handling is comparable between groups and RyR2cKO hearts do not display regional variability (**Figure 3A-B**). We analyzed the CaT parameters of each individual cell recorded in different regions of the heart. We found that CaT amplitude, time to peak and rise velocity were not different between groups, while 50% decay time was significantly prolonged in RyR2cKO myocytes (**Figure 3C-F**). Quantification of RyR2 expression in the same hearts using Western blots showed a reduction of >90% in RyR2cKO hearts (**Figure 3G-H**), consistent with our previous results. To evaluate whether RyR2 expression was uniformly reduced across RyR2cKO hearts we performed immunostaining in fixed slices from the same hearts used for Ca^2+^ imaging and Western blots (**Figure 4**). To identify the boundaries of individual cells, we co-stained for N-cadherin, a protein that localizes to the intercalated discs, and RyR2. RyR2 expression was robust in control hearts, and localization was in the expected striated pattern (**Figure 4A**). In contrast, RyR2cKO hearts showed almost no visible RyR2 signal and there were few interspersed myocytes with higher RyR2 signal than surrounding cells. Interestingly, N-cadherin intensity was slightly increased in RyR2cKO slices while DAPI intensity was decreased (**Figure 4C-D**). These data suggest that cardiomyocytes in RyR2cKO hearts have nearly normal Ca^2+^ dynamics despite an overall reduction of >90% in RyR2 expression and that CaTs do not originate only from cells that retain relatively higher or nearly normal RyR2 levels. Therefore, these results support our hypothesis that ECC possesses plasticity that can compensate for a dramatic homogeneous reduction in RyR2 expression.

**Figure 3.**
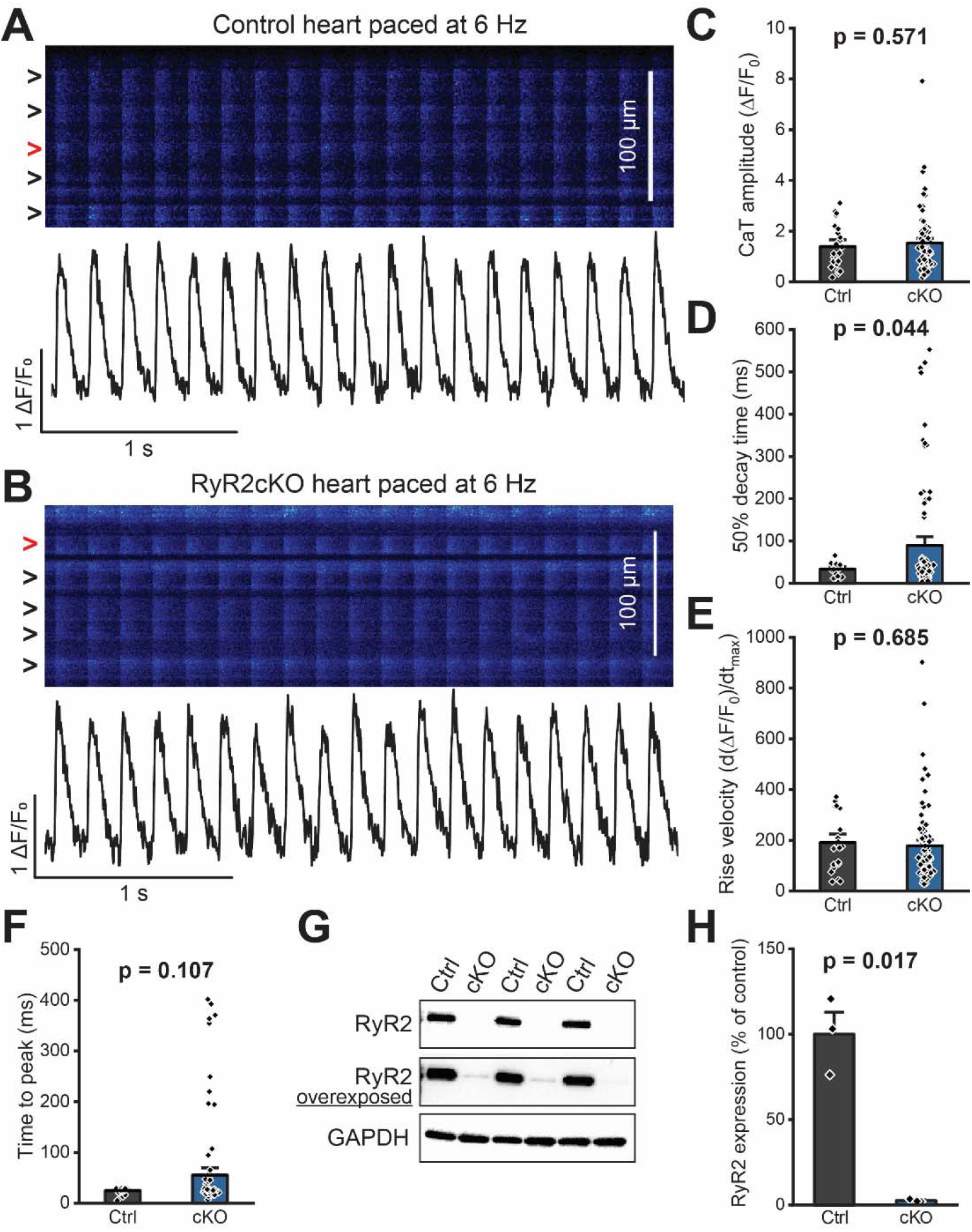
Whole-heart Ca^2+^ transients are indistinguishable between control and RyR2cKO hearts 3-weeks post-TAM. **A, B**) Representative line-scan imaging of Ca^2+^ transients (top) in control (Ctrl, A) and RyR2cKO (cKO, B) hearts loaded with Rhod-2 and perfused with normal Tyrode solution supplemented with blebbistatin and paced at 6 Hz. Ca^2+^ transient signal profile (bottom) from a representative cell (red arrow) of the recorded region. **C-F**) Quantification of CaT amplitude (C), 50% decay time (D), rise velocity (E), and time to peak (F) from individual cells as shown in panels A and B. N = 3 hearts per genotype; n = 23 Ctrl and 88 cKO cells. Linear mixed-effects model (LMM) with Bonferroni post-hoc. **G, H**) Representative Western blots and quantification of RyR2 expression in the same hearts used for Ca^2+^ imaging. GAPDH was used as loading control. N = 3 hearts per genotype. T-test.

**Figure 4.**
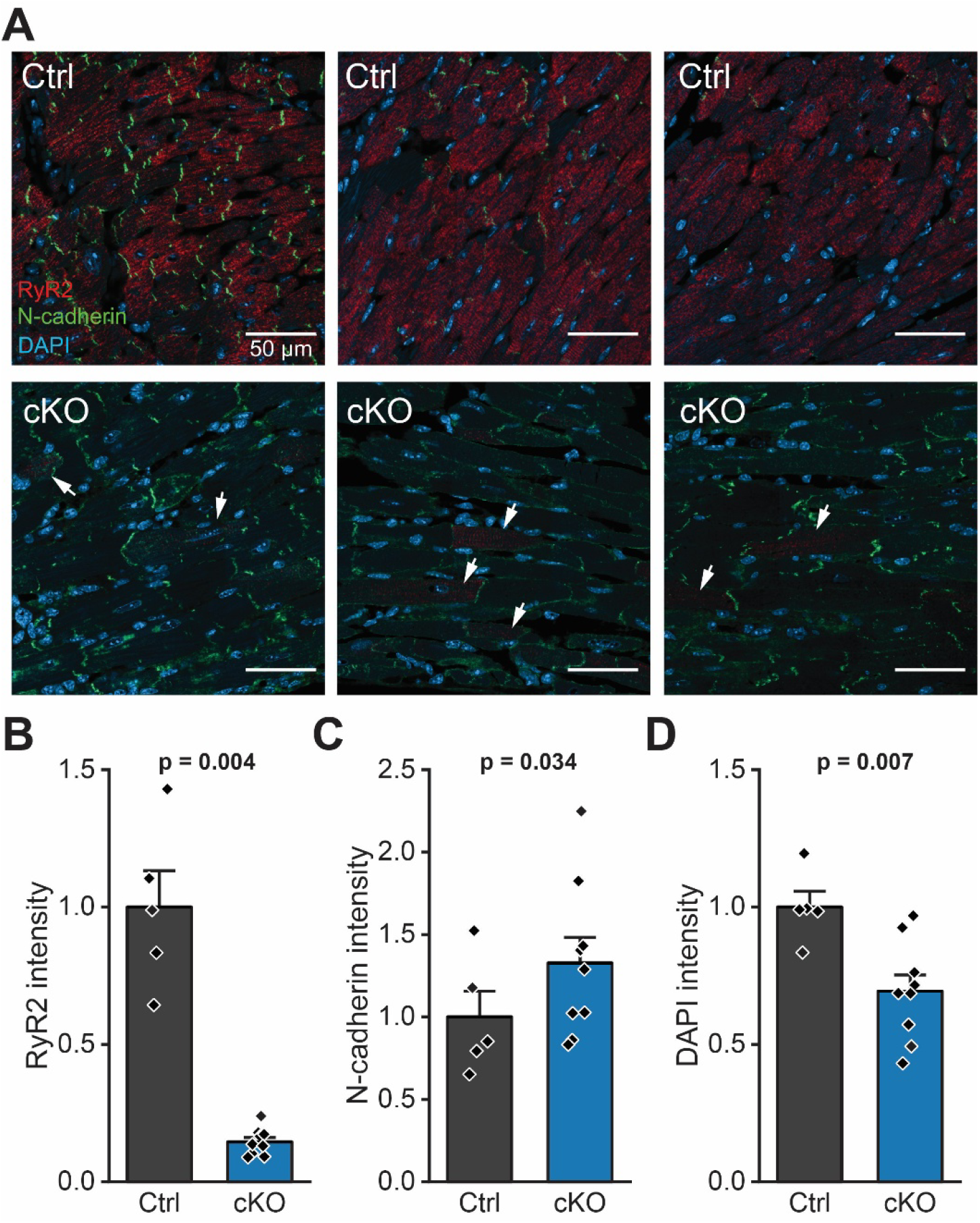
RyR2 expression is homogeneously depleted at the tissue level in RyR2cKO hearts. **A**) Representative images from tissue sections obtained from control (Ctrl) and RyR2cKO (cKO) fixed hearts stained for RyR2 (red), N-cadherin (green) and DAPI (blue). Hearts were collected 3-weeks after TAM injections. Scale bars are 50 µm long. All images were acquired and analyzed using the same parameters and settings for both control and RyR2cKO hearts. **B-D**) Quantification of RyR2 (B), N-cadherin (C), and DAPI (D) abundance as measured by fluorescence intensity normalized by the area of the image and normalized to the average signal in control hearts. Each data point represents a region imaged from a heart. N = 2 hearts with n = 5 regions for control, and N = 3 hearts with n = 8 regions for RyR2cKO. Linear mixed-effects model (LMM) and Bonferroni post-hoc.

### Single-cell Ca^2+^ handling is preserved in RyR2 knockout cardiomyocytes

To evaluate cellular Ca^2+^ dynamics at higher spatial and temporal resolution in the absence of cell-to-cell coupling, we measured CaTs in cardiomyocytes isolated from RyR2cKO and control mice 3 weeks post-TAM injection. Cells were paced with field stimulation at 1, 2, and 3 Hz, and CaT properties were analyzed (**Figure 5A**). CaTs at 1 Hz showed similar amplitude and decay time (**Figure 5B,C**); however, the kinetics of Ca^2+^ release were significantly slower in RyR2cKO myocytes, as shown by the reduced upstroke velocity and prolonged time to peak (**Figure 5D,E**). These changes were also present at 2 and 3 Hz stimulation frequencies **(Figure 6**). The SR Ca^2+^ load, measured during a pulse of caffeine at the end of the stimulation period, was significantly increased in RyR2cKO myocytes (**Figure 5F**), while fractional release was significantly decreased (**Figure 5G**). Hence, a >90% reduction in RyR2 expression does not overtly disrupt intracellular Ca^2+^ dynamics in cardiomyocytes.

**Figure 5.**
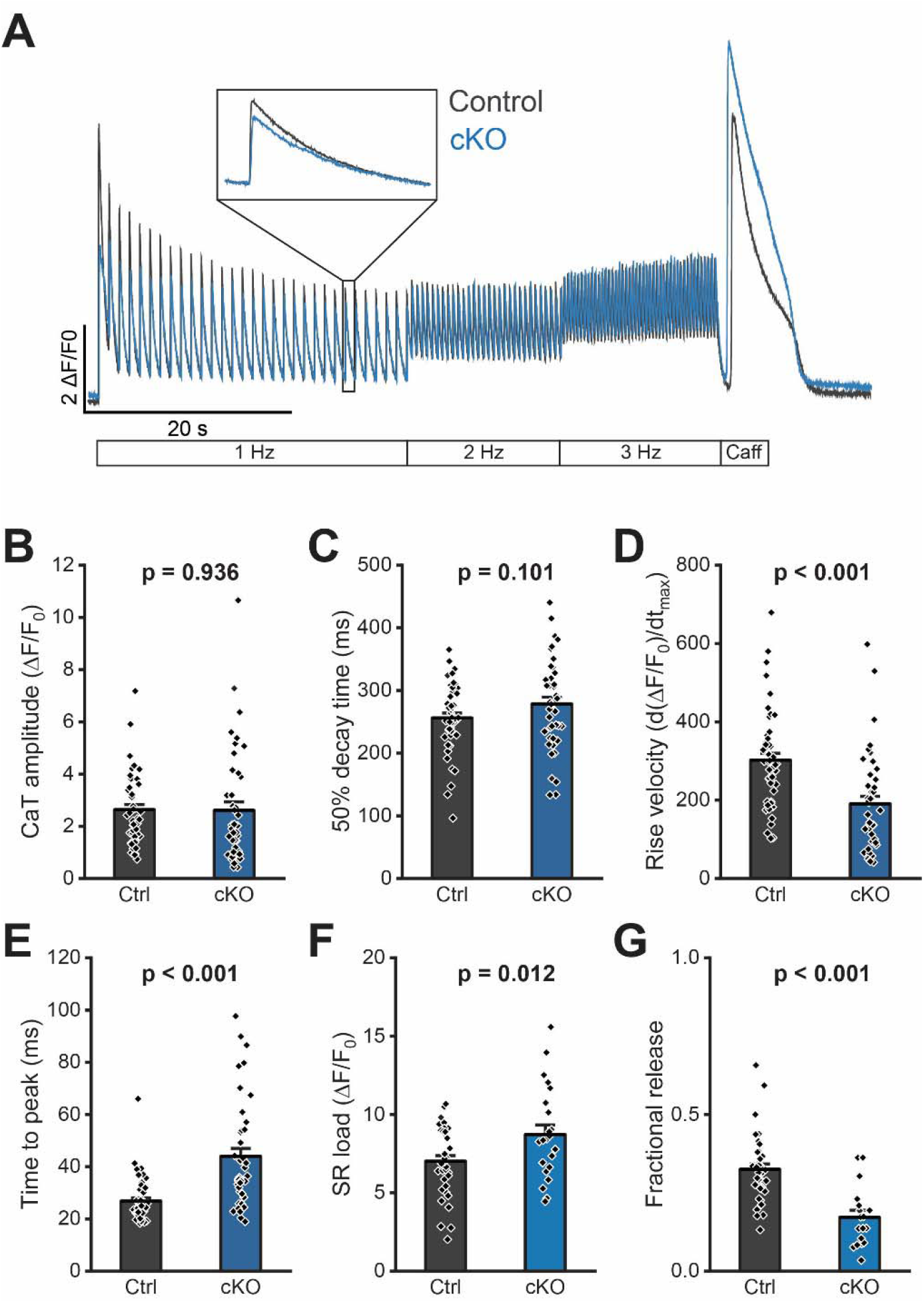
Ca^2+^ handling properties are comparable in control and RyR2cKO cardiomyocytes. **A**) Representative Ca^2+^ transient recordings in cardiomyocytes from control (Ctrl) and RyR2cKO (cKO) hearts at 3 weeks post-TAM injection and paced at 1, 2, and 3 Hz. The stimulation protocol is shown below traces. **B-E**) Quantification of CaT amplitude (B), 50% decay time (C), rise velocity (D), and time to peak (E) in Ctrl and cKO myocytes paced at 1 Hz. **F**) Sarcoplasmic reticulum (SR) load in control and cKO myocytes calculated during a pulse of caffeine (10 mM) at the end of the 3 Hz stimulation period. **G**) Fractional release estimated by dividing the CaT amplitude at 1 Hz by the SR load in control and cKO myocytes. N = 3 Ctrl and 4 cKO hearts; n = 48 Ctrl and 43 cKO myocytes for panels B-E. n = 38 for control and n = 23 cKO myocytes for panels F and G. Linear mixed-effects model (LMM) and Bonferroni post-hoc.

**Figure 6.**
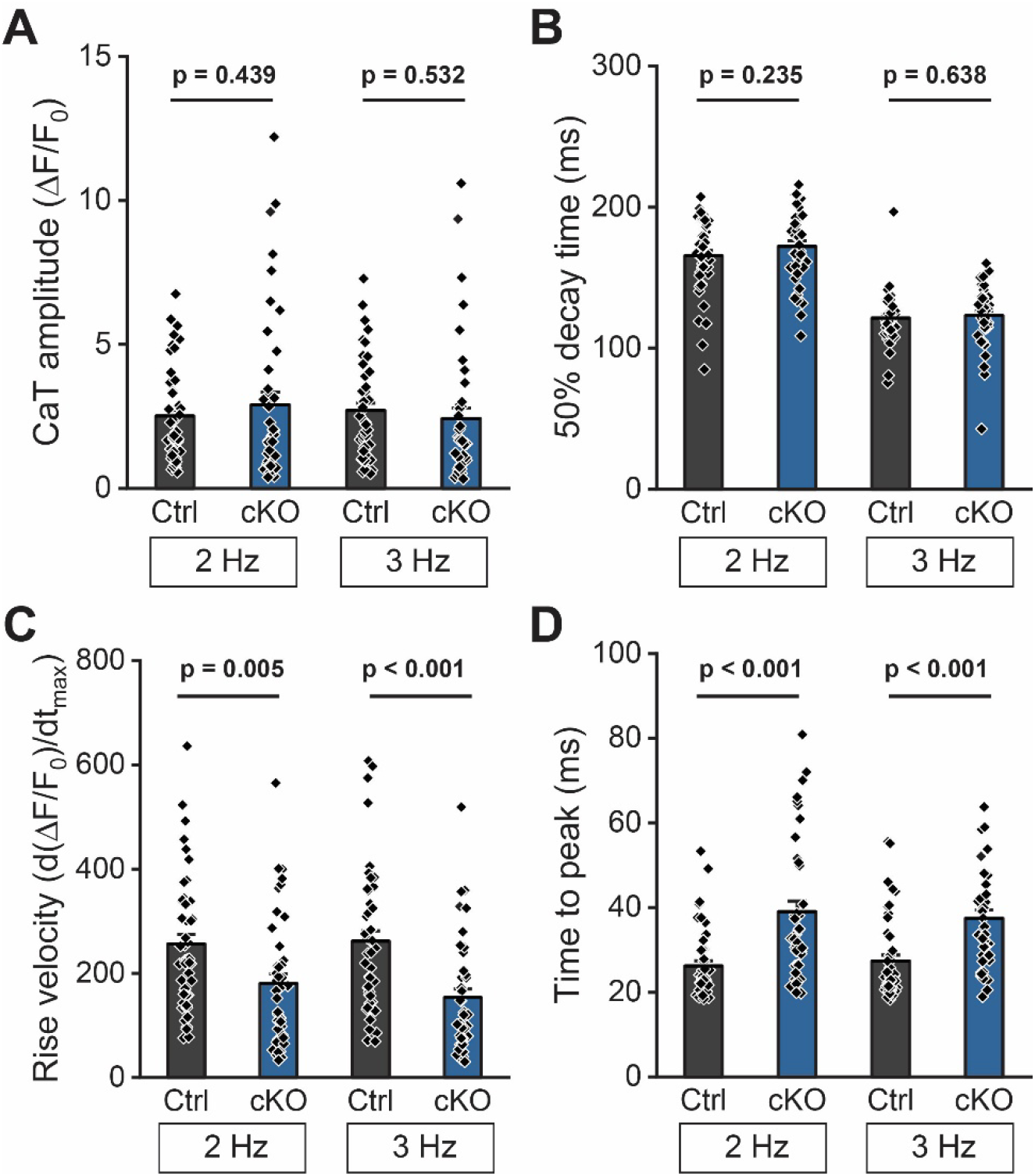
Ca^2+^ handling properties in control and RyR2cKO isolated cardiomyocytes at 2 and 3 Hz stimulation frequencies. **A-D**) CaT amplitude (A), 50% decay time (B), rise velocity (C), and time to peak (D) quantification in control and cKO myocytes measured from transients paced at 2 and 3 Hz, from same experiments shown in Figure 5. N = 3 Ctrl and 4 cKO hearts; n = 47 Ctrl and 42 cKO myocytes. Linear mixed-effects model (LMM) with Bonferroni post-hoc.

### Excitation-contraction coupling gain is preserved in RyR2cKO myocytes

CaT data showed that the few remaining RyR2 channels in RyR2cKO myocytes can preserve normal function in isolated cells. Nonetheless, changes in the coupling between Ca_V_1.2 and RyR2 can occur upon RyR2 depletion. To investigate this, we simultaneously measured *I*_Ca_ and the resulting CaT in isolated control and RyR2cKO myocytes. Representative *I*_Ca_ values across a range of voltages and the elicited CaTs are shown in **Figure 7A-B**. Coupling between Ca_V_1.2 and RyR2 was preserved across the different tested potentials in RyR2cKO myocytes. No significant changes in *I*_Ca_ and CaT amplitude were observed between groups, as shown in the current-voltage (**Figure 7C**) and fluorescence-voltage relationships (**Figure 7D**), respectively. We further calculated the ECC amplification factor or ECC gain by dividing the CaT amplitude by the trigger *I*_Ca_ at different testing potentials. ECC gain was nearly identical in control and RyR2cKO myocytes, with the exception of 40 mV, where there was a significant decrease in RyR2cKO myocytes (**Figure 7E**). Altogether, our results show that global Ca^2+^ handling can be preserved with low RyR2 expression. These results support the idea that sufficient Ca^2+^ release can be achieved by activation of a small number of RyR2 channels. Membrane capacitance (C_m_) measurements obtained during these experiments were also similar between groups control and RyR2cKO myocytes (187.5 ± 16.8 pF for control *versus* 190.1 ± 15.1 pF for RyR2cKO myocytes), suggesting the absence of cellular hypertrophy.

**Figure 7.**
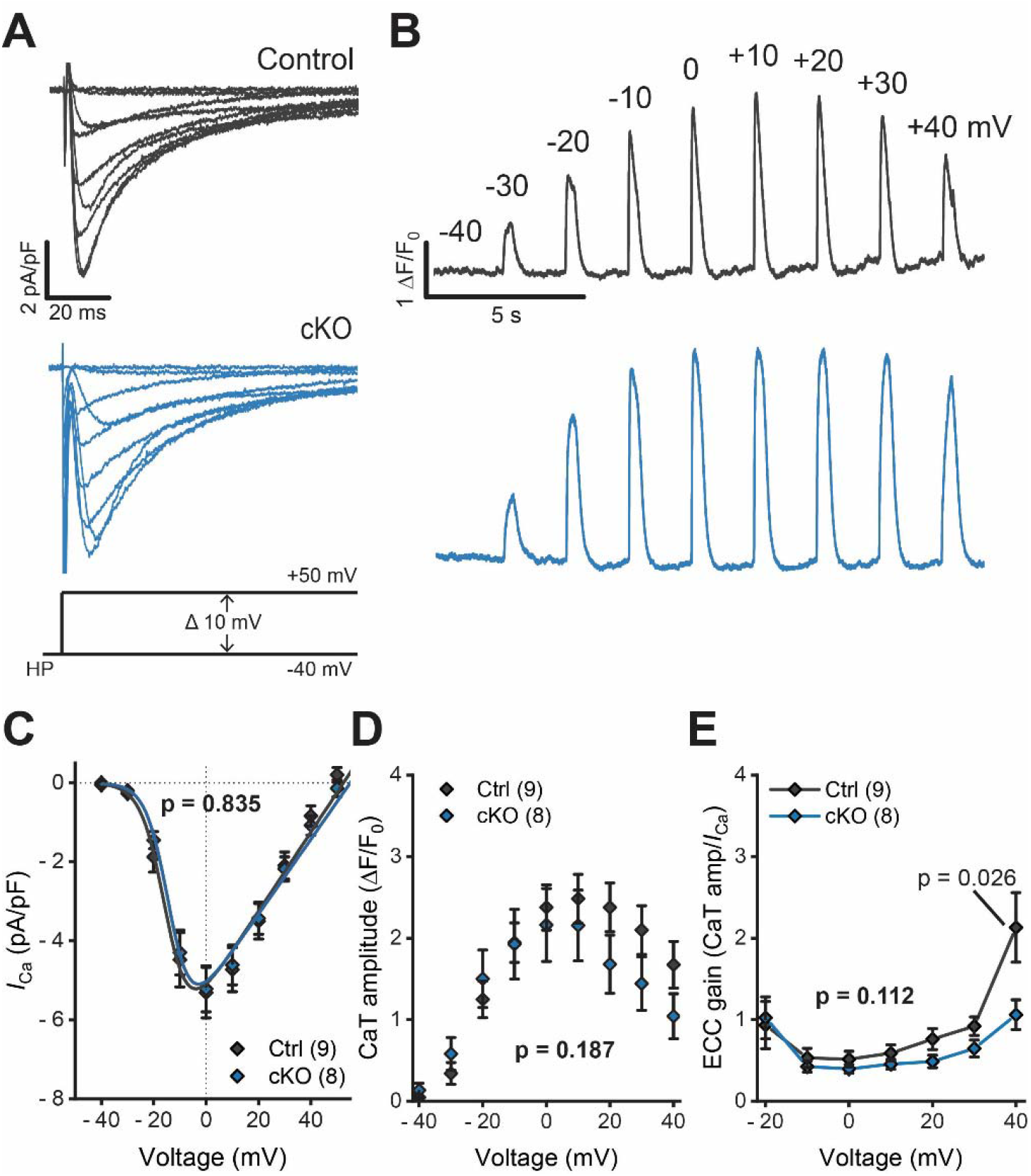
Excitation-contraction coupling gain is preserved under decreased RyR2 expression. **A, B**) Simultaneous recordings of L-type Ca^2+^ currents (*I*_CaL_, A) and Ca^2+^ transients (CaT, B) in control (Ctrl) and RyR2cKO (cKO) cardiomyocytes. The voltage step protocol is indicated at the bottom of the representative trace families; HP stands for holding potential. Voltage applied to elicit the different CaTs is indicated on top of the control representative trace for clarity. **C**) I-V curve from *I*_CaL_ control and cKO myocytes 3 weeks post-TAM. Solid lines show the Boltzmann fit for the experimental data. **D**) CaT amplitude elicited by Ca^2+^ current at the different test potentials in Ctrl and cKO myocytes. **E**) Excitation-contraction coupling gain in Ctrl and cKO myocytes, determined by dividing the CaT amplitude by its corresponding Ca^2+^ current. N = 3 mice per genotype. The number of cells specified in each panel. P-values shown in bold represent the comparison of the complete curve between groups. In panels C and D, no significant differences were observed at any voltage. For panel E, p-value pointing at the data point indicates a significant difference only for 40 mV. Linear mixed-effects model (LMM) with Repeated Measures and Bonferroni post-hoc.

To preserve ECC fidelity after RyR2 depletion, changes in the expression of other ECC proteins could compensate. To investigate this, we measured the levels of SERCA2, Ca_V_1.2, NCX and PLB expression in control and RyR2cKO hearts at three weeks post-TAM injection using Western blots. Interestingly, RyR2cKO hearts showed significantly reduced SERCA2 expression, while other ECC proteins were unchanged (**Figure 8A-E**).

**Figure 8.**
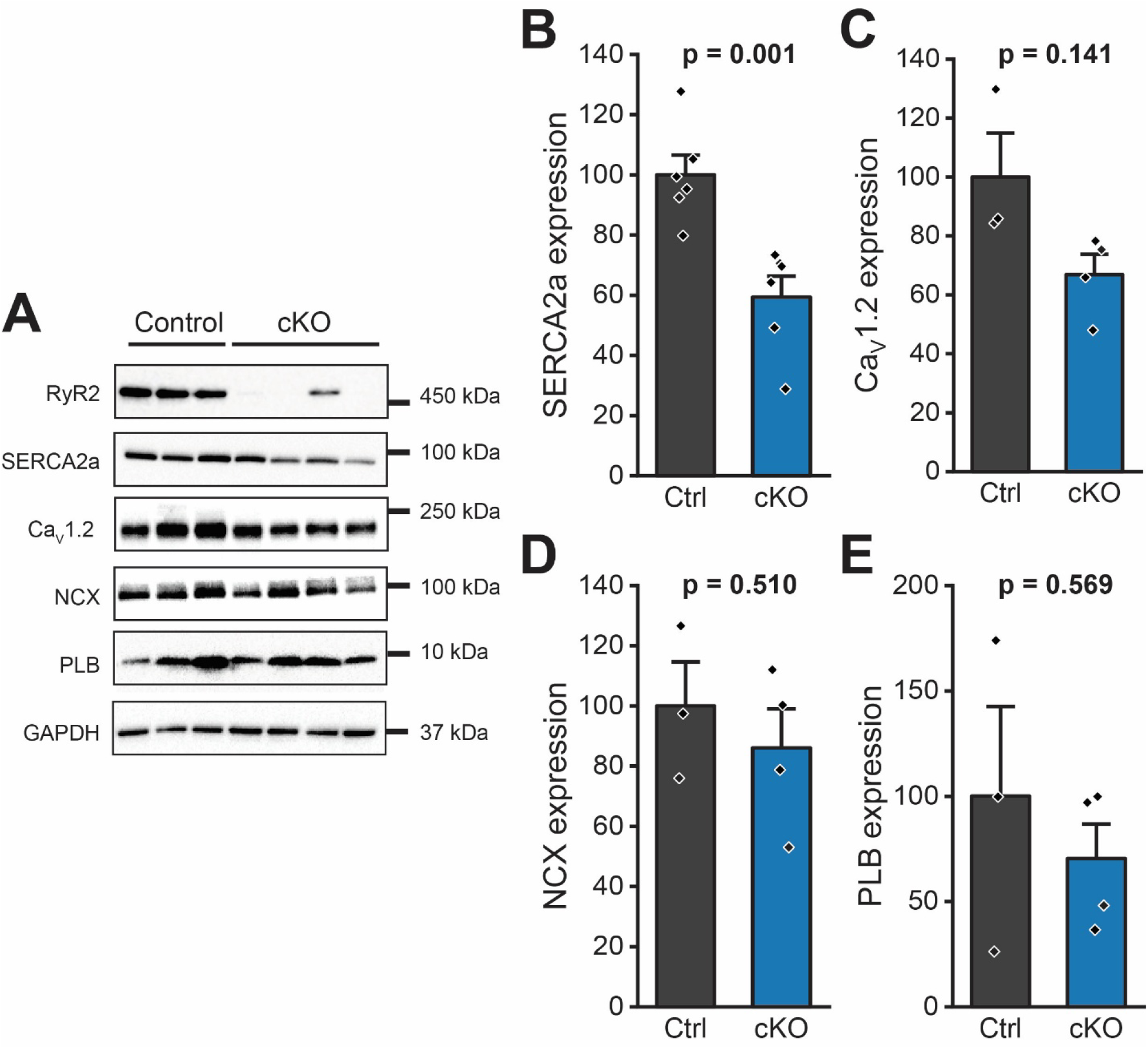
Excitation-contraction coupling protein expression shows minor changes in RyR2cKO hearts. **A**) Representative Western blots of relevant excitation-contraction coupling (ECC) proteins in control (Ctrl) and RyR2cKO (cKO) hearts at 3 weeks post-TAM injection. GAPDH was used as loading control. **B-E**) Quantification of SERCA2 (B), Ca_V_1.2 (C), NCX (D), and PLB (E) expression from Western blot analysis. All data were normalized to GAPDH and to the level of expression in control hearts. Data points represent one individual heart. RyR2, ryanodine receptor 2; SERCA2, sarco/endoplasmic reticulum Ca^2+^ ATPase; Ca_V_1.2, voltage-gated Ca^2+^ 1.2 channel; NCX, Na^+^/Ca^2+^ exchanger; PLB, phospholamban. N = 6 Ctrl and 6 cKO hearts. (Panel B); N = 3 Ctrl and 4 cKO hearts (Panels C-E). T-test.

### Other SR channels do not compensate for the loss of RyR2

RyR2 is one of multiple ion channels expressed in the SR. While other channels are not proposed to functionally interact with Ca_V_1.2 to promote ECC, is possible that they may contribute to compensate for the lower RyR2 expression. A recent study in human induced pluripotent stem cell-derived cardiomyocytes suggested that IP_3_ receptors (IP_3_Rs) can compensate for RyR2 deficiency (Starnes *et al*., 2024). To test this hypothesis, we measured IP_3_R expression in control and RyR2cKO hearts at three weeks post-TAM (**Figure 9A**). Western blot analysis showed a significant ∼20% reduction in IP_3_R expression in RyR2cKO hearts compared to controls (**Figure 9B**), suggesting that compensation is not through increased expression. Vacuole membrane protein 1 (VMP1) was recently reported as an SR Ca^2+^ leak channel that does not participate directly in ECC but may contribute to regulate SR load (Ma *et al*., 2026). Interestingly, Western blots showed that VMP1 expression was significantly reduced by ∼60% in RyR2cKO hearts compared to controls (**Figure 9C**). We hypothesize that VMP1 does not directly compensate for RyR2 depletion. Instead, reduced expression in RyR2cKO myocytes could constrain SR Ca^2+^ leak, contributing to increased SR content and Ca^2+^ release.

**Figure 9.**
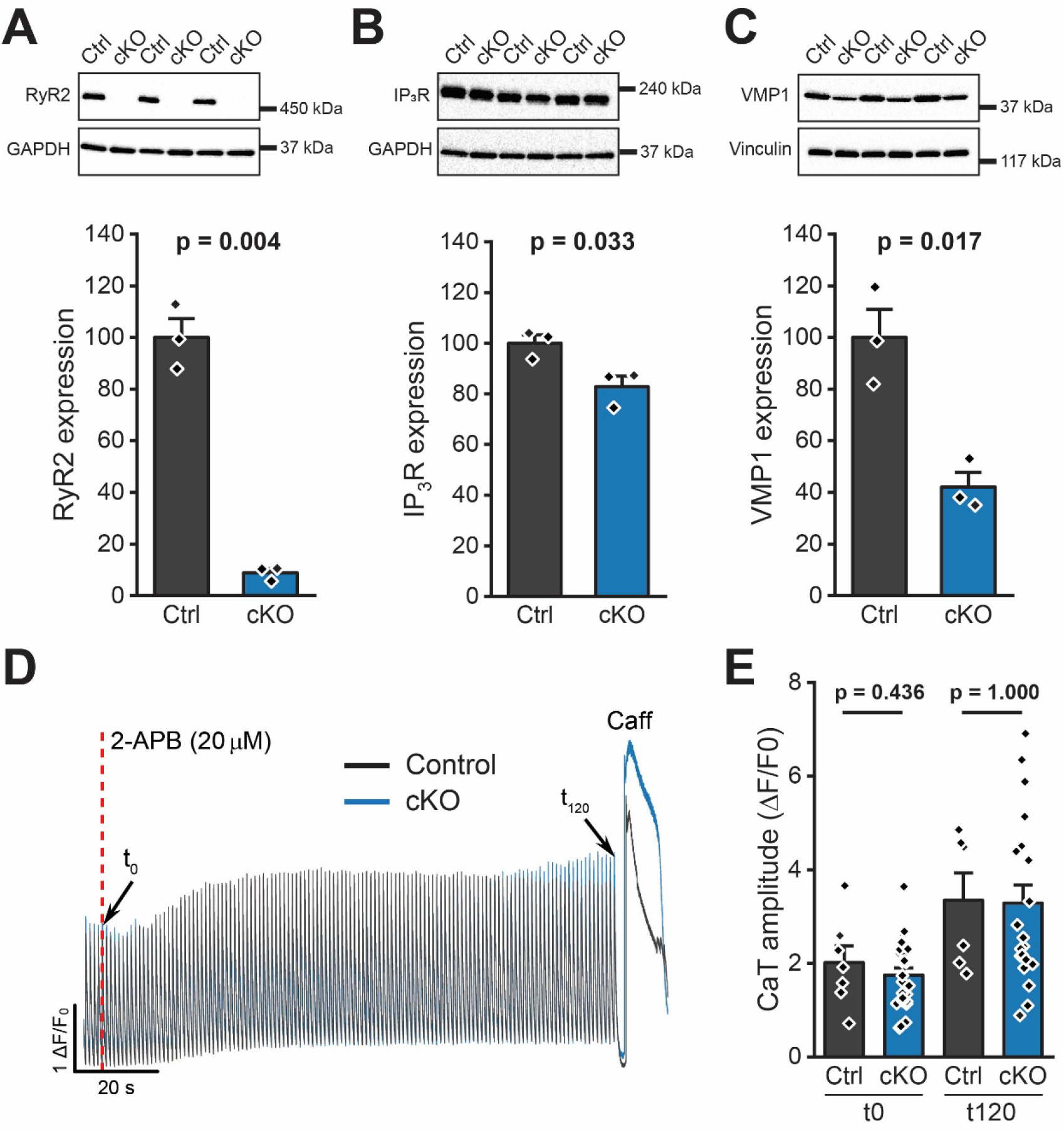
Compensation does not come from other SR Ca^2+^ release channels. **A-C**) Representative Western blots and quantification of the expression of RyR2 (A), IP_3_ receptor (IP_3_R) and vacuole membrane protein 1 (VMP1) in control (Ctrl) and RyR2cKO (cKO) hearts at 3 weeks post-TAM injection. GAPDH was used as loading control for RyR2 and IP_3_R, while vinculin was used as loading control for VMP1. N = 3 per genotype. T-test. **D**) Representative fluorescence profile of confocal CaTs in a control and a cKO cardiomyocyte at 3 weeks post-TAM during perfusion with 2-APB (20 µM, an IP_3_R blocker) over 2 min. **E**) CaT amplitude quantification at the start of 2-APB perfusion (t_0_) and after 2 min (t_120_). N = 1 Ctrl and 3 cKO hearts; n = 7 Ctrl and 24 cKO cardiomyocytes. Linear mixed-effects model (LMM) with Repeated Measures and Bonferroni post-hoc.

Finally, we recorded CaTs in RyR2cKO myocytes in the presence of 2-APB, an IP_3_R inhibitor (Maruyama *et al*., 1997). We hypothesized that CaT properties in RyR2cKO cells would be dramatically affected by 2-APB, potentially suppressing CaTs altogether, if IP_3_Rs provided functional compensation for the loss of RyR2. Cardiomyocytes were stimulated at 1 Hz for 30 s in normal Tyrode solution before perfusing 2-APB (20 µM) for 2 min (**Figure 9D**). 2-APB did not induce significant changes in CaT amplitude in control and RyR2cKO cells, although there was a tendency for increased CaT amplitude in both groups (**Figure 9E**). This suggests that in adult ventricular myocytes, IP_3_Rs do not compensate for the reduction in RyR2 expression.

## DISCUSSION

ECC is the fundamental physiological process that powers the heart. It relies on a complex network of proteins that sense membrane voltage, relay an intracellular signal that prompts Ca^2+^ release from the SR and recycle Ca^2+^ out of the cytosol. Despite the critical role of these players, we and others have shown that their expression can be reduced without overt cardiac dysfunction. For example, we previously demonstrated that rabbit hearts can withstand a ∼50% reduction in RyR2 expression free from a structural or functional phenotype (Zheng *et al*., 2022). Our new findings show that ECC is far more resilient to loss of RyR2 than previously appreciated. Despite a >90% reduction in RyR2 protein expression, global Ca²O handling, ECC gain, and whole-heart function remained remarkably preserved. Neither electro nor echocardiography detected electrical or structural remodeling, and both intact hearts and isolated myocytes maintained normal CaT amplitude. We only observed slower Ca^2+^ release kinetics, as demonstrated by the prolonged CaT time to peak, and elevated SR Ca²O load. These results demonstrate the functional capacity of the ECC apparatus to compensate for RyR2 expression without significant remodeling. The preservation of ECC fidelity without transcriptional remodeling and with minimal changes in other ECC proteins underscores a previously unrecognized form of homeostatic plasticity in the heart, positioning RyR2 abundance as a surprisingly flexible parameter rather than a strict determinant of ECC fidelity.

### RyR2 channel availability and clustering give ECC plasticity

We demonstrate that CaTs in intact hearts from control and RyR2cKO mice are indistinguishable even when RyR2 expression is below 10% of that of control hearts. This may seem like a surprising result; however, it is consistent with previous estimates. Bers considered the number of RyR2 channels expressed in cardiomyocytes (1.5-3.5 million channels), the total flux of SR Ca^2+^ required to increase intracellular Ca^2+^ to systolic levels (50 μmol/L cytosol), the mass of single Ca^2+^ sparks (7500 Ca^2+^ sparks at 40 fC/spark), and the conductance of RyR2 channels (2 fC/RyR2) to estimate that only ∼5% of RyR2 channels is needed to produce a normal contraction in basal conditions (Bers, 2001). Hence, even with less than 10% of RyR2 channels remaining in RyR2cKO myocytes, only half of those channels would be needed to produce a normal CaT. Ca^2+^ sparks are release events originating from localized RyR2 clusters, and the whole-cell CaT results from the temporal and spatial summation of sparks (Cheng & Lederer, 2008). Galice et al. previously demonstrated that approximately 10 RyR2 channels can produce nearly maximal Ca^2+^ spark amplitude, while cluster size controls the overall spontaneous spark frequency (Galice *et al*., 2018). More recently, Hou et al. showed that the mean open probability of RyR2 channels within a cluster at the peak of a Ca^2+^ spark is very low (∼0.42) and that all sparks are produced by the nearly simultaneous opening of 1 to 5 channels regardless of the size of the cluster or the distance between them (Hou *et al*., 2023). Therefore, macroscopic Ca^2+^ release estimates and Ca^2+^ spark analyses agree with our observations showing that Ca^2+^ cycling can be maintained with a significantly small fraction of the RyR2 channels normally available in cardiomyocytes. Hence, ECC plasticity may be achieved, at least in part, by the intrinsic biophysical properties of clustered RyR2 channels. Furthermore, there is an abundant RyR2 channel reserve, i.e., more channels available than cardiomyocytes require to produce a CaT under basal conditions.

### Autoregulation and other compensatory mechanisms

The idea that cardiomyocytes have an abundant RyR2 channel reserve that contributes to ECC fidelity is consistent with our results, but other compensatory mechanisms may also be in place. We first confirmed that *I*_Ca_ and ECC gain are preserved in RyR2cKO myocytes; hence, sarcolemmal Ca^2+^ fluxes do not compensate for RyR2 expression. Autoregulation of CaT dynamics was proposed by Eisner and colleagues as a passive mechanism that counteracts perturbations of RyR2 open probability by adjusting SR Ca^2+^ load and flux, thereby normalizing Ca^2+^ release (Eisner *et al*., 1998). Hence, pharmacological suppression of RyR2 function would reduce CaT amplitude only temporarily. The resulting buildup of Ca^2+^ in the SR would prompt larger release and normalize the CaT. A similar paradigm could apply when RyR2 expression is reduced. We previously reported that autoregulation was unlikely to compensate in a rabbit model with a constitutive decrease of ∼50% in RyR2. We concluded that chronic changes in RyR2 expression were fundamentally different from acute pharmacological inhibition of the channel; therefore, autoregulation could be superseded by other mechanisms such as the reorganization of RyR2 clusters (Zheng *et al*., 2022). Here, we confirm that this framework does not fully apply in the setting of acute depletion of RyR2. In RyR2cKO cardiomyocytes, we observed a normal CaT amplitude together with an increase in SR load and a reduction in SR fractional release (**Figure 5F,G**), suggesting that autoregulation is, indeed, balancing Ca^2+^ fluxes. Thus, SR Ca^2+^ content directly influences the magnitude of systolic release to preserve ‘normal’ Ca^2+^ homeostasis. This integrated view explains why RyR2 function has limited impact on steady-state contractility and instead highlights the trigger *I*_Ca_ and SR Ca^2+^ content as the central regulators of cardiac performance.

### Comparison of our results with previous studies

Several approaches have been taken to study the effects of reduced RyR2 expression in the heart. Constitutive and conditional knockout models have been reported with RyR2 levels ranging from 50 to 80% (Bround *et al*., 2012; Zhong *et al*., 2018; Zheng *et al*., 2022). While the focus and goals of each study vary, they consistently report that global Ca^2+^ handling and heart function remain unaffected in the setting of reduced RyR2 levels. Constitutive homozygous RyR2-KO is embryonic lethal in mice and rabbits (Takeshima *et al*., 1998; Zheng *et al*., 2022). Knocking out a single allele, constitutively or conditionally, reduces RyR2 protein expression by ∼50% and does not present cardiac functional or structural abnormalities (Zou *et al*., 2011; Bround *et al*., 2016; Zheng *et al*., 2022). We show that RyR2cKO mice can survive for several weeks without overt cardiac dysfunction (**Figure 1**). Mice eventually die, most likely when RyR2 is completely lost in a critical number of cardiomyocytes. Two studies reported that RyR2 depletion in adult mice was associated with the development of hypertrophy (Bround *et al*., 2012; Bround *et al*., 2013). In contrast, our echocardiography data show structurally normal RyR2cKO hearts (**Figure 2**) and membrane capacitance measurements isolated cardiomyocytes from control and RyR2cKO hearts suggested the absence of cellular hypertrophy. The discrepancies between previous studies and our results could be explained by the time points at which the measurements were conducted. Bround et al. reported transient hypertrophy in RyR2cKO mice four days post-TAM that was fully recovered by the eighth day. This is consistent with the known cardiac toxicity of the MerCreMer transgene used to induce Cre-lox recombination of the *Ryr2* gene. We used the same recombination system but purposely focused our studies at three weeks post-TAM, when the effect of TAM and MerCreMer activation waned. Nonetheless, we did not observe overt remodeling even at one week post-TAM. The absence of cardiac remodeling and preserved Ca^2+^ homeostasis at this timepoint, when RyR2 depletion exceeds 90%, underscores the plasticity of the ECC machinery.

### RyR2 and SERCA2 are coregulated after RyR2 knockout

ECC fidelity relies on the highly coordinated function of multiple proteins localized to the sarcolemma and the SR membrane. Interestingly, we found no significant changes in the expression of several major ECC players, including Ca_V_1.2, PLB and NCX. The notable exception was a significant decrease in the expression of SERCA2 in RyR2cKO hearts (**Figure 8**). This may seem counterintuitive, given that SR load was increased in RyR2cKO cardiomyocytes. Two mechanisms may contribute to this observation: autoregulation, as discussed earlier, and reduced VMP1 expression, which may limit non-RyR2-mediated SR Ca^2+^ leak (**Figure 9**) (Ma *et al*., 2026). Limiting Ca^2+^ reuptake can also constrain SR overload. Recent studies have shown that co-translational regulation of sarcolemmal ion channels modulates excitability in cardiomyocytes (Liu *et al*., 2016; Eichel *et al*., 2019). It is tantalizing to hypothesize that functionally related components of the ECC machinery may undergo a similar regulation mode. Remarkably, transcripts of SR proteins such as RyR2 and SERCA2 are locally synthesized to facilitate functional deployment, (Bogdanov *et al*., 2021) which could facilitate their co-translational interaction. Elucidating the mechanism by which knocking out RyR2 triggers a reduction in SERCA2, VMP1 and IP_3_R expression will require further investigation.

## Conclusion

In conclusion, we demonstrate that normal Ca^2+^ homeostasis and cardiac function can be maintained with RyR2 expression below 10% of the normal level. These findings demonstrate that ECC possesses intrinsic plasticity, as it quickly adapts to a significant loss of RyR2 to maintain normal function. This plasticity is achieved through a combination of three concomitant mechanisms. (1) Cardiomyocytes have a large RyR2 reserve, and only a fraction of the channels expressed are needed to sustain ECC in basal conditions. This may not be applicable to stress conditions. Indeed, we previously observed dyssynchronous Ca^2+^ release in cardiomyocytes under adrenergic stress when RyR2 levels are reduced to ∼50% (Zheng *et al*., 2022). (2) Autoregulation, which maintains SR Ca^2+^ release by balancing fluxes in conditions where RyR2 function is acutely modified. (3) Co-regulation of key proteins, including SERCA2 and VMP1, which is likely to contribute to maintaining SR Ca^2+^ content within a workable range that enhances autoregulation while preventing arrhythmogenic overloading.

### Pathophysiological relevance

Our study challenges the idea that reduced RyR2 expression, as seen in disease, is a pathological mechanism. In different cardiac disorders, post-translational modifications in RyR2 promote arrhythmogenic Ca^2+^ leak (Dridi *et al*., 2020). Instead, reducing channel expression could limit the deleterious effect of dysfunctional RyR2s and limit arrhythmogenesis. Gain-of-function (GoF) and loss-of-function (LoF) mutations in RyR2 are linked to inherited arrhythmia syndromes (Priori *et al*., 2002; Sun *et al*., 2021). Our data prompt an important distinction between the functional outcomes of LoF mutations and a reduction in channel expression. RyR2 tetramers heterogeneously assembled from wild-type and mutant subunits are dysfunctional, as previously noted (Loaiza *et al*., 2013; Zhao *et al*., 2015). In contrast, homogeneous reduction of wild-type RyR2 channel expression does not result in loss of function. Hence, RyR2 can bridge the electrical signal into contraction even at low expression levels. This intrinsic ECC plasticity opens the possibility to target mutant RyR2 channels for degradation in heterozygous patients (maintaining a single WT allele) while preserving normal function, as recently proposed (Moore *et al*., 2024).

## DATA AVAILABILITY STATEMENT

All data supporting the findings of this study are available upon reasonable request.

## AUTHOR CONTRIBUTIONS

FJA conceived the project. FJA and EBRP conceptualized most experiments and wrote the manuscript. EBRP collected and analyzed most of the data and prepared the figures. AMG performed and analyzed Western blots. DWK analyzed echocardiography data. ICG and SE performed the CHX chase assay. All authors discussed the results and approved the manuscript.

## ACKNOWLEDGEMENTS

We thank Ashley Weichmann from the Small Animal Imaging and Radiotherapy Facility (SAIRF) and Magdaline Baus from the Translational Research Initiatives in Pathology (TRIP) laboratory for their help with echocardiography and tissue immunohistochemistry, respectively.

